# bModelTest: Bayesian phylogenetic site model averaging and model comparison

**DOI:** 10.1101/020792

**Authors:** Remco R Bouckaert, Alexei J Drummond

**Affiliations:** Centre for Computational Evolution, University of Auckland, Auckland, NZ.; Department of Computer Science, University of Auckland, Auckland, NZ.; Max Planck Institute for the Science of Human History, Jena, Germany.

**Keywords:** Model averaging, Model selection, Model comparison, Statistical phylogenetics, ModelTest, Phylogenetic model averaging, Phylogenetic model comparison, Substitution model, Site model

## Abstract

**Background:** Reconstructing phylogenies through Bayesian methods has many benefits, which include providing a mathematically sound framework, providing realistic estimates of uncertainty and being able to incorporate different sources of information based on formal principles. Bayesian phylogenetic analyses are popular for interpreting nucleotide sequence data, however for such studies one needs to specify a site model and associated substitution model. Often, the parameters of the site model is of no interest and an ad-hoc or additional likelihood based analysis is used to select a single site model.

**Results:** bModelTest allows for a Bayesian approach to inferring and marginalizing site models in a phylogenetic analysis. It is based on trans-dimensional Markov chain Monte Carlo (MCMC) proposals that allow switching between substitution models as well as estimating the posterior probability for gamma-distributed rate heterogeneity a proportion of invariable sites and unequal base frequencies. The model can be used with the full set of time-reversible models on nucleotides, but we also introduce and demonstrate the use of two subsets of time-reversible substitution models.

**Conclusion:** With the new method the site model can be inferred (and marginalized) during the MCMC analysis and does not need to be pre-determined, as is now often the case in practice, by likelihood-based methods. The method is implemented in the bModelTest package of the popular BEAST 2 software, which is open source, licensed under the GNU Lesser General Public License and allows joint site model and tree inference under a wide range of models.

## Background

One of the choices that needs to be made when performing a Bayesian phylogenetic analysis is which site model to use. A common approach is to use a likelihood-based method like ModelTest [1], jModelTest [2], or jModelTest2 [3] to determine the site *model*. The site model is comprised of (i) a substitution model defining the relative rates of different classes of substitutions and (ii) a model of rate heterogeneity across sites which may include a gamma distribution [4] and/or a proportion of invariable sites [5, 6]. The site model recommended by such likelihood-based method is then often used in a subsequent Bayesian phylogenetic analysis. This analysis framework introduces a certain circularity, as the original model selection step requires a phylogeny, which is usually estimated by a simplistic approach. Also, by forcing the subsequent Bayesian phylogenetic analysis to condition on the selected site model, the uncertainty in the site model can’t be incorporated into the uncertainty in the phylogenetic posterior distribution. A more statistically rigorous and elegant method is to co-estimate the site model and the phylogeny in a single Bayesian analysis, thus alleviating these issues.

One way to select substitution models for nucleotide sequences is to use reversible jump between all possible reversible models [7], or just a nested set of models [8]. An alternative is to use stochastic Bayesian variable selection [9], though this does not address whether to use gamma rate heterogeneity or invariable sites. Wu et al. [10] use reversible jump for substitution models and furthermore select for each site whether to use gamma rate heterogeneity or not. Since the method divides sites among a set of substitution models, it does not address invariable sites, and only considers a very limited set of five (K80, F81, HKY85, TN93, and GTR) substitution models.

We introduce a method which combines model averaging over substitution models with model averaging of the parameters governing rate heterogeneity across sites using reversible jump. Whether one considers the method to be selecting the site model, or averaging over (marginalizing over) site models depends on which random variables are viewed as parameters of interest and which are viewed as nuisance parameters. If the phylogeny is viewed as the parameter of interest, then bModelTest provides estimates of the phylogeny averaged over site models. Alternatively if the site model is of interest, then bModelTest can be used to select the site model averaged over phylogenies. These are matters of post-processing of the MCMC output, and it is also possible to consider the interaction of phylogeny and site models. For example one could construct phylogeny estimates conditional on different features of the site model from the results of a single MCMC analysis.

The method is implemented in the bModelTest package of BEAST 2 [11] with GUI support for BEAUti making it easy to use. It is open source and available under LGPL licence. Source code, installation instructions and documentation can be found at https://github.com/BEAST2-Dev/bModelTest.

## Implementation

All time-reversible nucleotide models can be represented by a 4 × 4 instantaneous rate matrix:

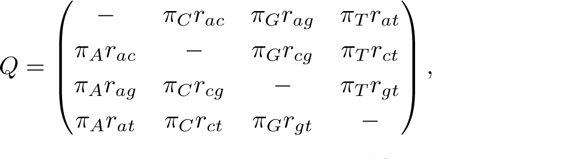

with six rate parameters *r*_*ac*_, *r*_*ag*_, *r*_*at*_, *r*_*cg*_, *r*_*ct*_ and *r*_*gt*_ and four parameters describing the equilibrium base frequencies Π = (*π*_*A*_, *π*_*C*_, *π*_*G*_, *π*_*T*_). A particular restriction on the rate parameters can conveniently be represented by a six figure model number where each of the six numbers corresponds to one of the six rates in the alphabetic order listed above. Rates that are constrained to be the same, have the same integer at their positions in the model number. For example, model 123456 corresponds to a model where all rates are independent, named the general time reversible (GTR) model [12]. Model 121121 corresponds to the HKY model [13] in which rates form two groups labelled transversions (1 : *r*_*ac*_ = *r*_*at*_ = *r*_*cg*_ = *r*_*gt*_) and transitions (2 : *r*_*ag*_ = *r*_*ct*_). By convention, the lowest possible number representing a model is used, so even though 646646 and 212212 represent HKY, we only use 121121.

There are 203 reversible models in total [7]. However, it is well known that transitions (A↔C, and G↔T substitutions) are more likely than transversions (the other substitutions) [14, 15]. Hence grouping transition rates with transversion rates is often not appropriate and these rates should be treated differently. We can restrict the set of substitution models that allow grouping only within transitions and within transversions, with the exception of model 111111, where all rates are grouped. This reduces the 203 models to 31 models (see Figure 1 and details in Appendix). Alternatively, if one is interested in using named models, we can restrict further to include only Jukes Cantor [16, 17] (111111), HKY [13] (121121), TN93 [18] (121131), K81 [19] (123321), TIM [20] (123341), TVM [20] (123421),and GTR [12] (123456). However, to facilitate stepping between TIM and GTR during the MCMC (see proposals below) we like to use nested models, and models 123345 and 123451 provide intermediates between TIM and GTR, leaving us with a set of 9 models (Figure 1).

**Figure 1.**
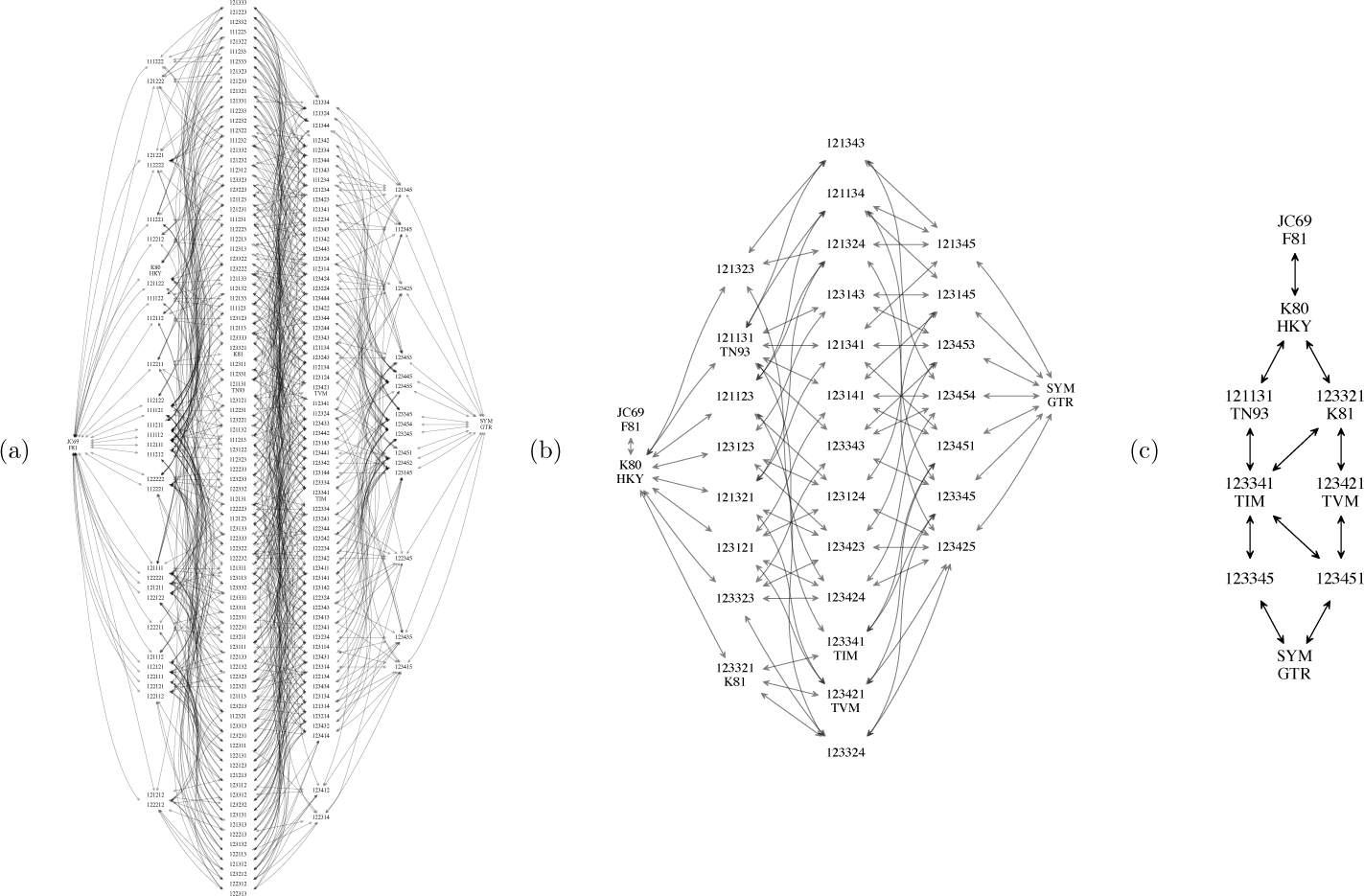
Model spaces. The model spaces supported by bModelTest. (a) All reversible models, (b) transition/transversion split models, and (c) named models. Arrows indicate which models can be reached by splitting a model. Note all models with the same number of groupings are at the same height.

The state space consists of the following parameters:

- the model number *M*,
- a variable size rate parameter (depending on model number) *R*,
- a binary variable to indicate whether 1 or *k* > 1 non-zero rate categories should be used,
- a shape parameter *α*, used for gamma rate heterogeneity when there are *k* > 1 rate categories,
- a binary variable to indicate whether or not a category for invariable sites should be used,
- the proportion of invariable sites *p*_*inv*_,

Rates *r*_*ac*_, *r*_*ag*_, *r*_at_, *r*_cg_, *r*_*ct*_ and *r*_*gt*_ are determined from the model number M and rate parameter *R*. Further, we restrict *R* such that the sum of the six rates 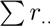 equals 6 in order to ensure identifiability. This is implemented by starting each rate with value 1, and ensuring proposals keep the sum of rates in (see details on proposals below).

### Prior

By default, bModelTest uses the flat Dirichlet prior on rates from [7]. From empirical studies [14, 15], we know that transition rates tend to be higher than transversion rates. It makes sense to encode this information in our prior and bModelTest allows for rates to get a different prior on transition rates (default log normal with mean 1 and standard deviation of 1.25 for the log rates) and transversion rates (default exponential with mean 1 for the rates).

An obvious choice for the prior on models is to use a uniform prior over all valid models. As Figure 1 shows, there are many more models with 3 parameters than with 1. An alternative allowed in bModelTest is to use a uniform prior on the number of parameters in the model. In that case, Jukes Cantor and GTR get a prior probability of 1/6, since these are the only models with 0 and 5 degrees of freedom respectively. Depending on the model set, a much lower probability is assigned to each of the individual models such that the total prior probability summed over models with *K* parameters, *p*(*K*) = 1/6 for *K* ∈ {0,1, 2, 3, 4, 5}.

For frequencies a Dirichlet(4,4,4,4) prior is used, reflecting our believe that frequencies over nucleotides tend to be fairly evenly distributed, but allowing a 2.2% chance for a frequency to be under 0.05. For *p*_*inv*_ a Beta(4,1) prior on the interval (0,1) is used giving a mean of 0.2 and for *α* an exponential with a mean 1. These priors only affect the posterior when the respective binary indicator is 1.

### MCMC proposals

The probability of acceptance of a (possibly trans-dimensional) proposal [21] is

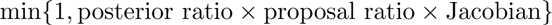

where the posterior ratio is the posterior of the proposed state *S’* divided by that of the current state S, the proposal ratio the probability of moving from *S* to *S’* divided by the probability of moving back from *S’* to *S*, and the Jacobian is the determinant of the matrix of partial derivatives of the parameters in the proposed state with respect to that of the current state [21].

#### Model merge/split proposal

For splitting (or merging) substitution models, suppose we start with a model *M*. To determine the proposed model *M’*, we randomly select one of the child (or parent) nodes in the graph (as shown in Figure 1). This is in contrast to the approach of Huelsenbeck *et al* [7], in which first a group is randomly selected, then a subgrouping is randomly created. For any set of substitution models organised in an adjacency graph our merge/split operator applies, making our graph-based method easier to generalise to other model sets (e.g. the one used in [22]). If there are no candidates to split (that is, model *M* = 123456 is GTR) the proposal returns the current state (this proposal is important to guarantee uniform sampling of models). Likewise, when attempting to merge model *M* = 111111, the current state is proposed (*M’* = 111111). Let *r* be the rate of the group to be split. We have to generate two rates *r*_*i*_ and *r*_*j*_ for the split into groups of size *n*_*i*_ and *n*_*j*_. To ensure rates sum to 6, we select *u* uniformly from the interval (−*n*_*i*_*r*, *n*_*j*_*r*) and set *r*_*i*_ = *r* + *u*/*n*_*i*_ and *r*_*j*_ = *r* − *u*/*n*_*j*_.

For a merge proposal, the rate of the merged group r from two split groups *i* and *j* with sizes *n*_*i*_ and *n*_*j*_, as well as rates *r*_*i*_ and *r*_*j*_ is calculated as 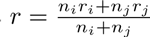.

When we select merge and split moves with equal probability, the proposal ratio for splitting becomes

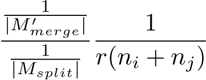

where 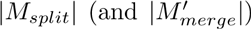 is the number of possible candidates to split (and merge) into from model *M* (and *M’* respectively). The proposal ratio for merging is

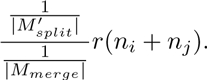

The Jacobian for splitting is 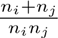 and for merging it is 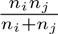

#### Rate exchange proposal

The rate exchange proposal randomly selects two groups, and exchanges a random amount such that the condition that all six rates sum to 6 is met. A random number is selected from the interval [0, *δ*] where *δ* is a tuning parameter of the proposal (*δ* is automatically optimized to achieve the desired acceptance probability for the data during the MCMC chain). Let *n*_*i*_, *r*_*i*_, *n*_*j*_ and *r*_*j*_ as before, then the new rates are 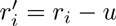 and 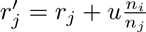. The proposal fails when 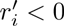.

The proposal ratio as well as the Jacobian are 1.

#### Birth/death proposal

Birth and death proposals set or unset the category count flag and sample a new value for *α* from the prior when the flag is set. The proposal ratio is *d*(*α’*) for birth and 1/*d*(*α*) for death where *d*(.) is the density used to sample from (by default an exponential density with a mean of 1).

Likewise for setting the indicator flag to include a proportion of invariable sites and sampling *p*_*inv*_ from the prior. The Jacobian is 1 for all these proposals.

#### Scale proposal

For the *α*, we use the standard scale operator in BEAST 2 [11], adapted so it only samples if the category count flag is set for α. Likewise, for *plnv* this scale operator is used, but only if the indicator flag to include a proportion of invariable sites is set.

## Results and Discussion

Since implementation of the split/merge and rate exchange proposals is not straightforward, nor is derivation of the proposal ratio and Jacobian, unit tests were written to guarantee their correctness and lack of bias in proposals (available on **https://github.com/BEAST2-Dev/bModelTest**).

To validate the method we performed a simulation study by drawing site models from the prior, then used these models to generate sequence data of 10K sites length on a tree (in Newick (A:0.2,(B:0.15,C:0.15):0.05)) with three taxa under a strict clock. The data was analysed using a Yule tree prior, a strict clock and bModelTest as site model with uniform prior over models and exponential with mean one for transversions and log-normal with mean one and variance 1.25 for transition rates. A hundred alignments were generated with gamma rate heterogeneity and a hundred without rate heterogeneity using a BEASTShell [23] script. Invariant sites can be generated in the process and are left in the alignment.

Comparing the model used to generate the alignments with inferred models is best done by comparing the individual rates of these models. Figure 2 shows the rate estimates for the six rates against the rates used to generate the data. Clearly, there is a high correlation between the estimated rates and the ones used to generate (R^2^ > 0.99 for all rates). Results were similar with and without rate heterogeneity. Note values for rates AG and CT (middle panels) tend to be higher than the transversion rates due to the prior they are drawn from.

**Figure 2.**
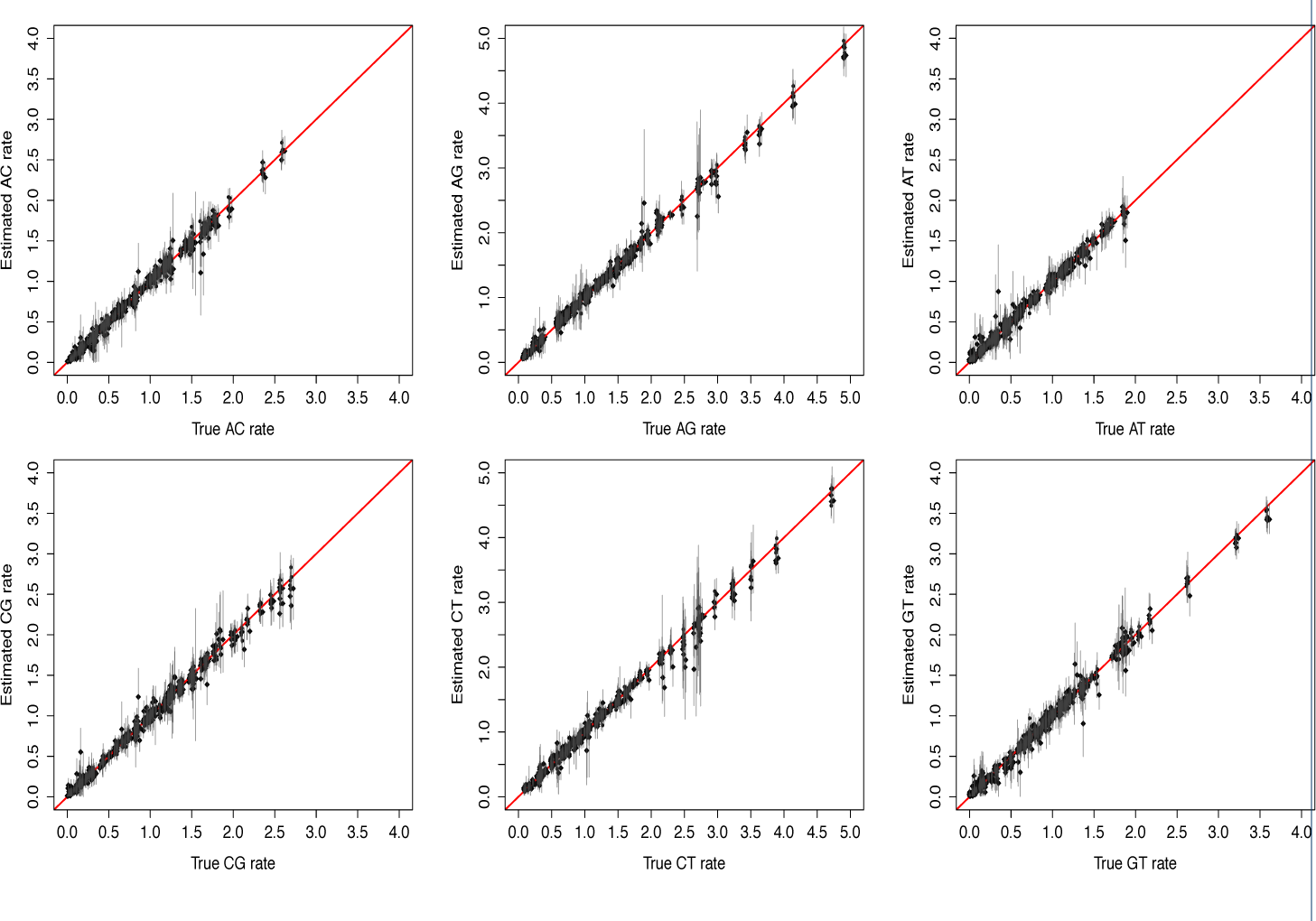
Accuracy of estimated substitution rates. True rates (horizontal) against estimated rates (vertical) in simulated data for 3 taxa. In reading order, rate AC, AG, AT, CG, CT and GT. Diamonds are for estimates when no rate heterogeneity was used to simulate the data, circles are for estimates with rate heterogeneity. Error bars represent 95% HPD intervals for each estimate.

Table 1 summarises coverage of the various parameters in the model, which is defined as the number of experiments where the 95% HPD of the parameter estimate contains the value of the parameter used to generate the data. The rows in the table show the four different models of rate heterogeneity among sites; plain means a single category without gamma or invariable sites, +G for discrete gamma rate categories, +I for two categories, one being invariable, and +G+I for discrete gamma rate categories and one invariable category. Furthermore, the experiment was run estimating whether base frequencies were equal or not. The first four rows are for data simulated with equal frequencies, the latter four with unequal frequencies. The last row shows results averaged over all 800 experiments. On average, one would expect the coverage to be 95% if simulations are drawn from the prior [24], so each entry in Table 1 has an expected value of 95, but can deviate due to small sample size. According to the binomial probability distribution there is a ~ 1.1% chance of seeing 89 or less successes when sampling 100 times with a success rate of 0.95. The sample size for the mean rows is 800, so is expected to be much closer to 95%.

**Table 1.**
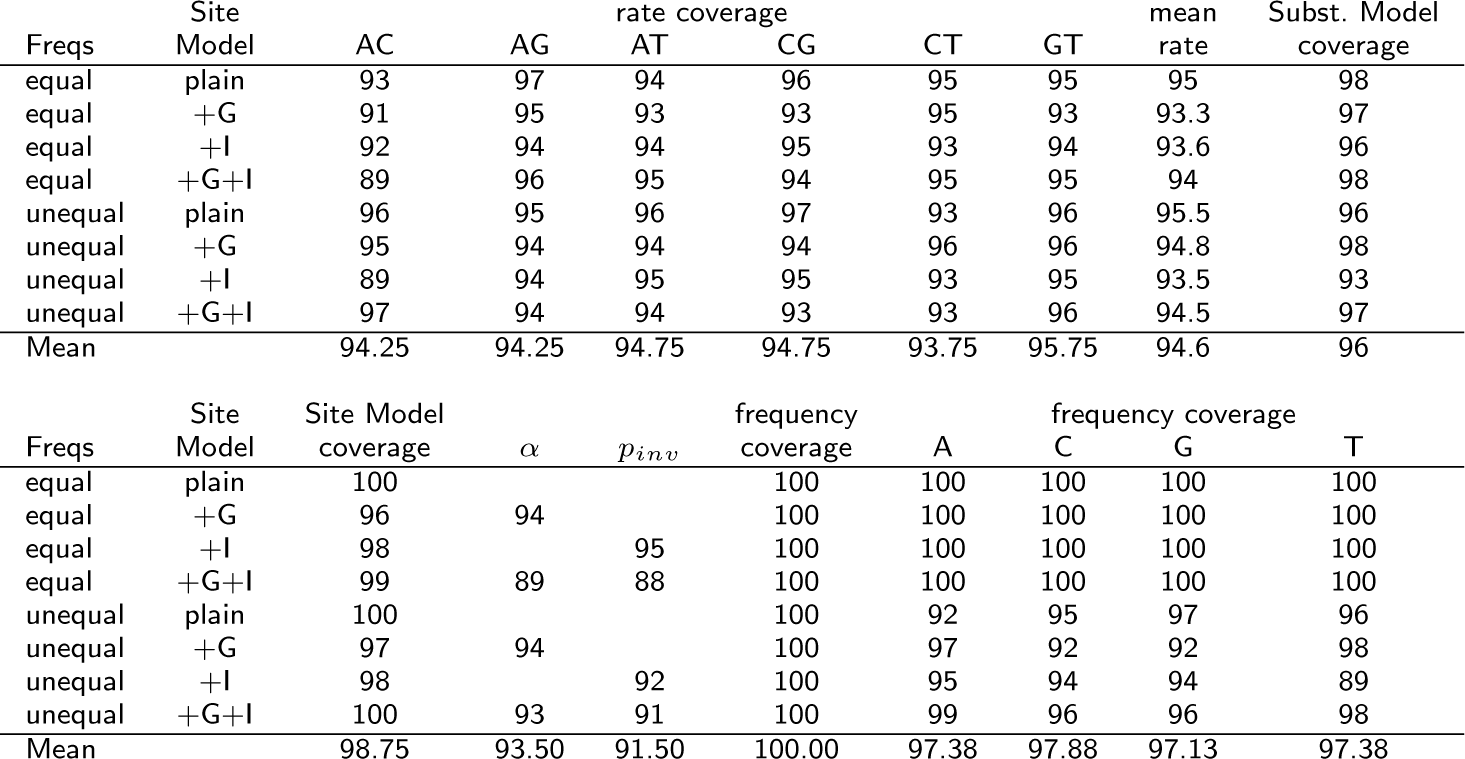
Coverage summary for simulation study. The first column lists the frequency and site models used to generate the data, and the last row is the mean coverage over all 800 runs. Coverage for rate parameters and frequencies is defined as the number of replicate simulations in which the true parameter value was contained in the estimated 95% HPD interval. The mean rate column contains the coverage averaged over all six rate coverage columns (i.e. the proportion of the 600 parameter estimates whose values were contained in their respective 95% HPD intervals. For details of substitution model coverage see text. The site model coverage is the number of replicate simulations that contained the correct model specification for rate heterogeneity across sites in the 95% credible set of models. Columns *α* and *pinv* are coverages of the shape and proportion invariable parameter *conditioned on sampling from the true site model*.

Coverage of rate estimates and frequencies are as expected, as shown in the table. Substitution model coverage is measured by first creating the 95% credible set of models for each simulation and then counting how often the model used to generate the data was part of the 95% credible set. The 95% credible set is the smallest set of models having total posterior probability ≥ 0.95. As Table 1 shows, model coverage is as expected (Subst. Model coverage column). The situation with gamma shape parameter estimates and proportion of invariable sites is not as straightforward as for the relative rates of the substitution process. The site model coverage can be measured in a similar fashion: the site model coverage column shows how often the 95% credible sets for the four different site models (plain, +G. +I and +G+I) contains the true model used to generate the data. The coverage is as expected. When looking at how well the shape parameter (*α* column in Table 1) and the proportion invariable sites (*p*_*inv*_ column in the table) is estimated, we calculated the 95% HPD intervals for that part of the trace where the true site model was sampled. Coverage is as expected when only gamma rate heterogeneity is used, or when only a proportion of invariable sites is used, but when both are used an interaction between the two site model categories appears to slightly reduce the coverage of both parameters. In these experiments the coverage for the frequency estimates for the individual nucleotides was as expected. In summary, the statistical performance of the model is as expected for almost all parameters except for the case where gamma and a proportion of invariable sites are used due to their interaction as discussed further below.

To investigate robustness of the approach, we repeated the study with a log normal uncorrelated relaxed clock [25] with a gamma(*α* = 30, *β* = 0.005) prior over the standard deviation for the log normal distribution. Trees with 5 taxa were randomly sampled from a Yule prior with log normal distribution (the birth rate was drawn from a distribution with a mean of the rate of 5.5, and a standard-deviation of the log-rate of 0.048) giving trees with mean height ≈ 0.25 and 95% HPD interval of 0.015 to 0.7. The study as outlined above was repeated, and results are summarised in Table S1, which looks very similar to that of Table 1. So, we conclude that the model is not sensitive to small variation in molecular clock rates among branches.

Figure 3 shows histograms of estimated posterior probability of gamma-distributed rate heterogeneity across sites for the data sets simulated over 5 taxa. When data was generated without gamma-distributed rate heterogeneity across sites, the posterior probability was often estimated to be close to zero (left of Figure 3), while the posterior probability was estimated to be close to one for most of the analyses on data in which gamma rate heterogeneity was present (middle of Figure 3).^[1]^ When rate heterogeneity was present, shape estimates were fairly close to the ones used to generate the data (right of Figure 3). However, there were quite a few outliers, especially when the shape parameter was high (although this is harder to see on a log-log plot which was used here because of the uneven distribution of true values).

**Figure 3.**
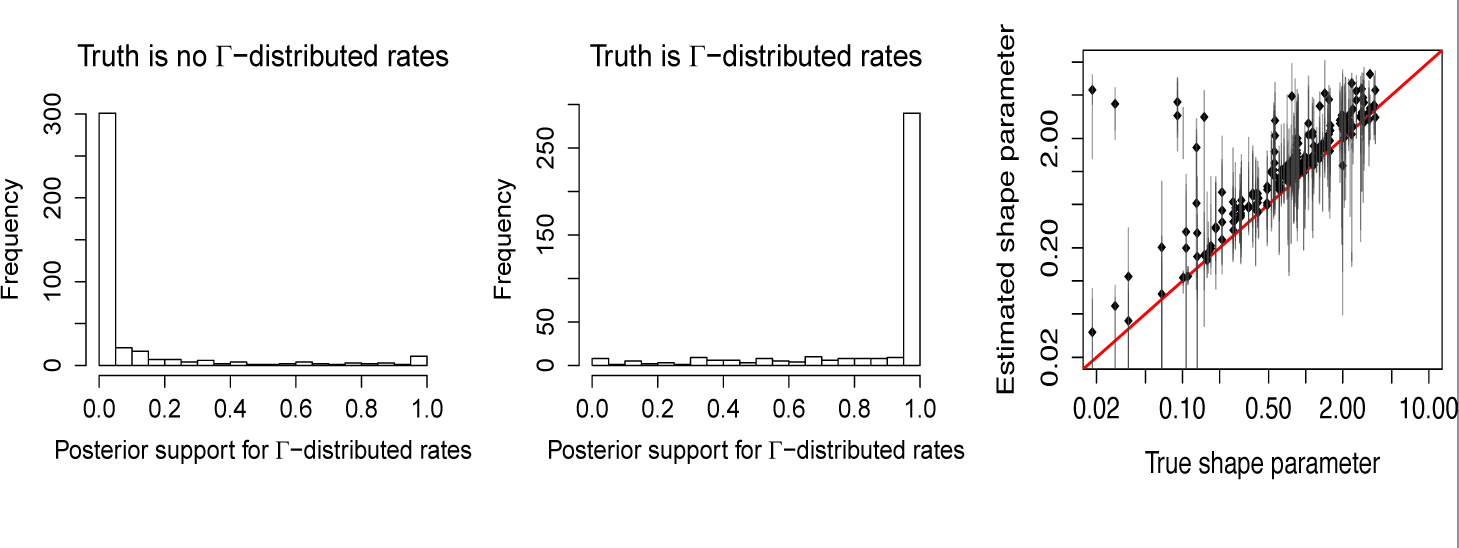
Accuracy of inference of rate heterogeneity across sites. Posterior probability for inclusion of gamma rate heterogeneity when the data is generated without (left) and with (middle) rate heterogeneity for 5 taxa. Right, True gamma shape parameter (horizontal) against estimated shape parameter (vertical) when rate heterogeneity is used to generate the data.

This can happen due to the fact that when the gamma shape is small, a large proportion of sites gets a very low rate, and may be invariant, so that the invariable category can model those instances. The mean number of invariant sites was 6083 when no rate heterogeneity was used, while it was 6907 when rate heterogeneity was used, a difference of about 8% of the sites.

Figure 4 shows similar plots as Figure 3 but for the proportion of invariable sites for 5 taxa.^[2]^ Empirically for the parameters that we used for our simulations, it appears that if there are less than 60% invariant sites, adding a category to model them does not give a much better fit. When a proportion of invariable sites was included in the simulation, there was a high correlation between the true proportion and the estimated proportion of invariable sites.

**Figure 4.**
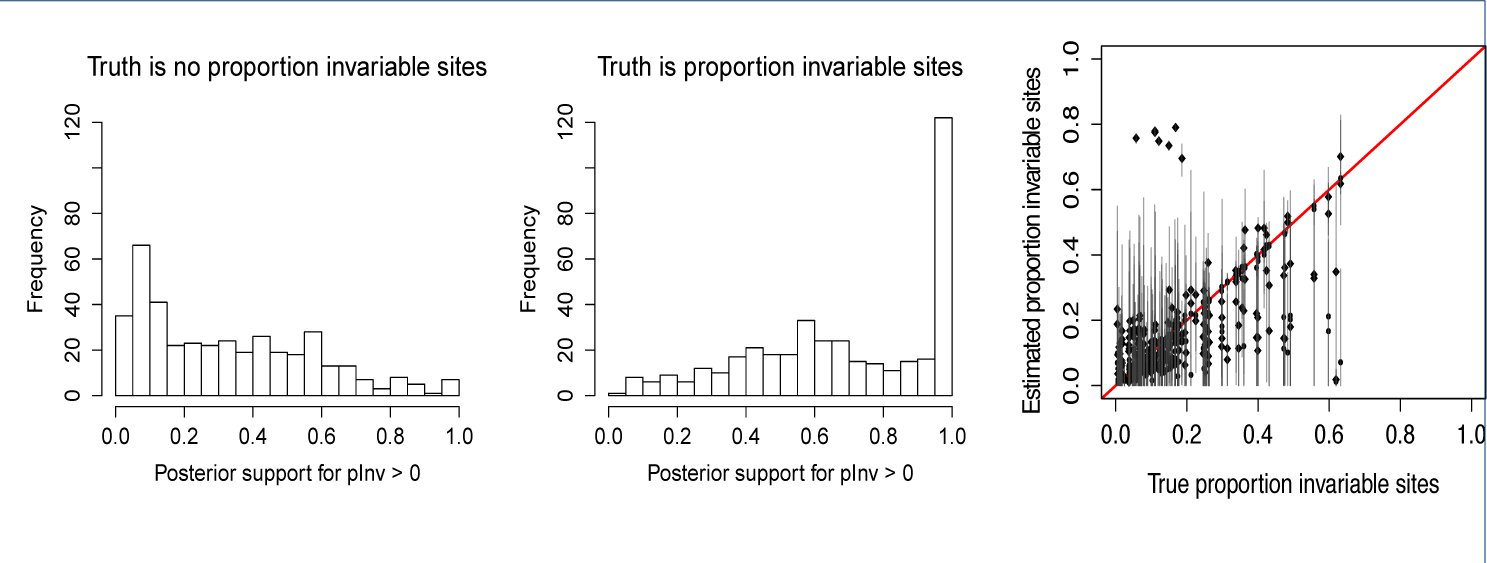
Accuracy of inference of proportion of invariant sites. Posterior probability for inclusion of a proportion of invariant sites when the data is generated without (left) and with (middle) invariant sites for 5 taxa. Right, empirical proportion invariant in alignment (horizontal) against estimated proportion of invariant sites (vertical) when a proportion invariable category is used to generate the data.

The same study with 5 taxa was repeated with the substitution model fixed to HKY and GTR, but estimating the other parts of the model. Results are summarised in Tables S2 and S3 respectively. Fixing the model to HKY results in severe degradation of accuracy in all parameter and model estimates. The lack of coverage of frequency estimates when the true model has equal frequencies suggests that lack of degrees of freedom in substitution model parameters is compensated by estimating frequencies instead of keeping them equal. So substitution model misspecification can result in considerable misspecification of the remainder of the model. Results when fixing the substitution model to GTR shows a table with results very similar to that of bModelTest, however the substitution model parameters have on average a 95% HPD interval of size 0.17 while that of bModelTest is only 0.13. The extra parameters that need to be estimated for GTR compared to bModelTest result in more uncertain estimates, and thus more uncertainty in the analysis.

To see the impact of the model set, the experiment was repeated with sampling from all 203 reversible models instead of using only the 30 transition/transversion split models. Results are shown in Table S4, which do not differ substantially from Table 1. Further, to investigate the effect of the number of taxa and sequence length, the study was repeated with 16 taxa and sequence lengths 1K and 0.5K base pairs long under a relaxed clock as before. Results are summarised in Tables S5 and S6 respectively. The tables do not show significant differences to Table 1 or degradation with decreasing sequence length, so the ability of our Bayesian method to correctly estimate the posterior distribution of substitution models and their parameters does not appear to depend substantially on sequence length or number of taxa.

### Comparison with jModelTest

We ran jModelTest version 2.1.10 [3] on the sequence data used for the last simulation study with 5 taxa (using all reversible models, since only that set is the same for both jModelTest and bModelTest) and the two simulation studies using 16 taxa and compared the substitution model coverage (with settings -BIC -AIC -f -g 4 -i -s 203). For each dataset, we collected the top models according to the AIC and BIC criteria such that the cumulative weight exceeded 95% of the models as shown in the jModelTest output and registered whether the true model was contained in the resulting set. Results are summarised in Table S7, which shows that both AIC and BIC do not cover the true model 95% of the time as would be desirable. For some combinations the coverage is close to the desirable value (88.4% for AIC with 5 taxa) and for some it is much lower (60.1% for BIC with 0.5K length sequences and 16 taxa). Coverage of both AIC and BIC appears to decrease with increasing number of taxa and decreasing sequence length in contrast to bModelTest, which has coverage of ~ 95% for all scenarios. jModelTest uses a single maximum likelihood tree and it seems that increasing uncertainty in the true tree (by increasing the number of taxa or decreasing sequence length) results in an increasing chance of incorrect model weights from jModelTest.

In practice, users of Bayesian phylogenetic packages only use the most highly weighted model returned by jModelTest. Table S7 shows how often the best fitting model according to AIC and BIC matches the true model, which ranges from 72.6% for BIC on 5 taxa to 29.9% for AIC on 0.5K length sequences and 16 taxa, suggesting that the probability of model misspecification using this approach increases with phylogenetic uncertainty.

To compare the application of bModelTest to jModelTest (with settings −f −i −g 4 −s 11 −AIC −a) we applied both to two real datasets. The first data set used was an alignment from 12 primate species [26] (available from BEAST 2 as file examples/nexus/Primates.nex) containing 898 sites. In this case the model recommended by jModelTest was TPM2uf+G and the substitution model TPM2 (=121323) has the highest posterior probability using bModelTest (21.12% see Appendix for full list of supported models) when empirical frequencies are used. However, when frequencies are allowed to be estimated, HKY has highest posterior probability (16.19%), while TPM2 (10.25%) has less posterior probability then model 121123 (14.09%). So, using a maximum likelihood approach (jModelTest and/or empirical frequencies) makes a substantial difference in the substitution model being preferred. Figure 5 left shows the posterior probabilities for all models, and it shows that the 95% credible set is quite large for the primate data. Figure 5 middle and 5 right show correlation between substitution model rates. The former shows correlation between transversion rate AC (horizontally) and transition rate AG (vertically). One would not expect much correlation between these rates since the model coverage image shows there is little support for these rates to be shared. However, since HKY is supported to a large extent and the rates are constrained to sum to 6, any proposed change in a transition rate requires an opposite change in transversion rates in order for the sum to remain 6. So, when sampling HKY, there is a linear relation between transition and transversion rates, which faintly shows up in the Figure 5 (middle). Figure 5 (right) shows the correlation between transversion rates AC and AT. Since they are close to each other, a large proportion of the time rate AC and AT are linked, which shows up as a dense set of points on the AC=AT line.

**Figure 5.**
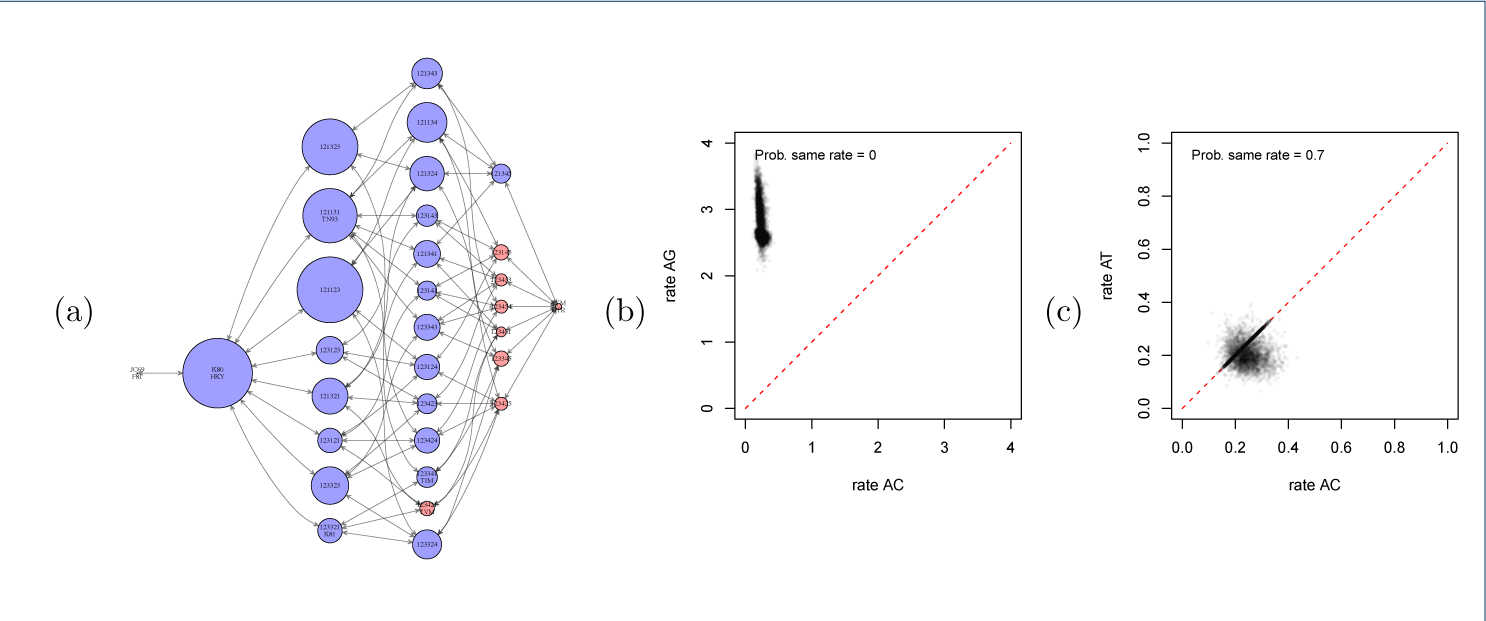
Posterior inference on primate data. Model distribution for primate data using the transition/transversion split models (left). Numbers on x-axis correspond to models in Appendix. The middle panel plots rates *A* ↔ *C* versus *A* ↔ *G* (middle) and the right panel plots *A* ↔ *C* versus *A* ↔ *T*.

The second data set used was an alignment of 31 sequences of 9030 sites of coding hepatitis C virus (HCV) from [27]. It was split into two partitions, the first containing codon 1 and 2 positions (6020 sites) and the second all codon 3 positions (3010 sites). Figure 6 left show the model distributions for the first partition at the top and second at the bottom. The 95% credible sets contain just 7 and 6 models respectively, much smaller than those for the primate data as one would expect from using longer, more informative sequences. Note that the models preferred for the first partition have transition parameters split while for the second partition models where partitions are shared have higher posterior probability, resulting in quite distinct model coverage images. For the first partition, bModelTest recommends TIM2+I+G. TIM2 is model 121343, the model with highest posterior probability according to bModelTest, as shown in Figure 6. For the second partition, jModelTest recommends GTR+G, and though GTR is in the 95% credible set, it has a lower posterior probability than TVM, even though TVM was considered by jModelTest. Again, we see a substantial difference in likelihood and Bayesian approaches. The correlation between transition rates *A* ↔ *G* and *C* ↔ *T* as well as between two transversion rates *A* ↔ *C* and *A* ↔ *T* are shown in Figures 6 top middle and right for the first partition and Figures 6 bottom middle and right for the second. The transition rates *A* ↔ *G* and *C* ↔ *T* have a posterior probability of being the same of 0.024 in the first partition, whereas the posterior probability is 0.66 in the second partition containing only 3rd positions of the codons. This leads to most models for the first partition distinguishing between *A* ↔ *G* and *C* ↔ *T*, while for the second partition most models share these rates. For the two transversion rates *A* ↔ *C* and *A* ↔ *T* the partitions display the opposite relationship, with the second partition preferring to distinguish them. As a result, overall the two partitions only have one model in common in their respective 95% credible sets, but that model (GTR) has quite low posterior probability from both partitions.

**Figure 6.**
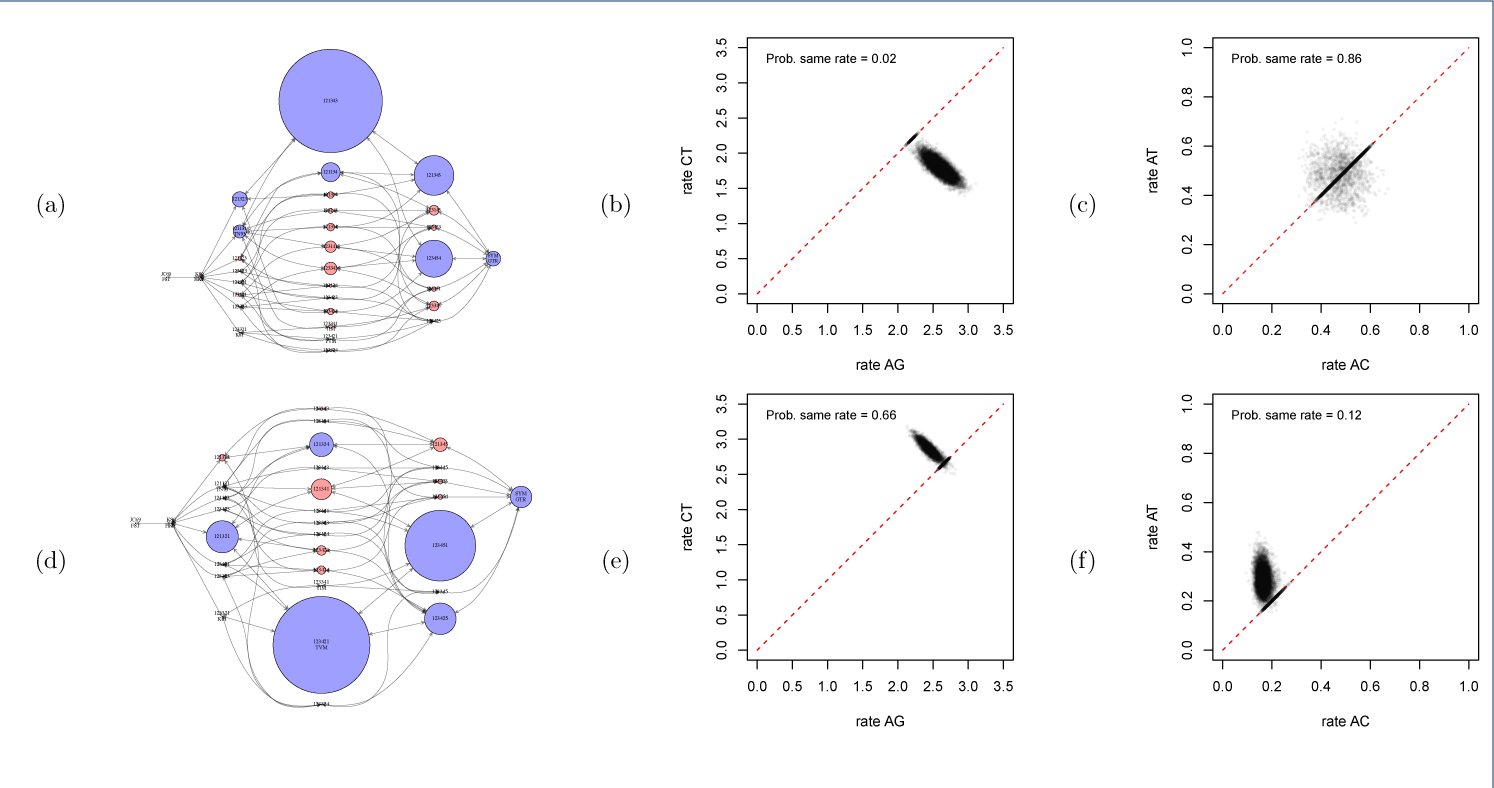
Posterior inference on HCV data. Like Figure 5, but the data is split into two partitions, the first containing codon positions 1+2 (panel a, b and c) and second containing codon position 3 (panel d, e and f). The partitions support distinctly different site models. The left panels show the posterior distribution over models, the middle panel plots transition rates *A* ↔ *G* versus *C* ↔ *T*, and the right panel plots transversion rates *A* ↔ *C* versus *A* ↔ *T*.

### Implementation details

The calculation of the tree likelihood typically consumes the bulk (≫ 90%) of computational time. Note that for a category with invariable sites, the rate is zero, hence only sites that are invariant (allowing for missing data) contribute to the tree likelihood. The contribution is 1 for those sites for any tree and for any parameter setting, so by counting the number of invariant sites, the tree likelihood can be calculated in constant time. Switching between with and without gamma rate heterogeneity means switching between one and k rate categories, which requires *k* time as much calculation. Having two tree likelihood objects, one for each of these two scenarios, and a switch object that selects the one required allows use of the BEAST 2 updating mechanism [9] so that only the tree likelihood that needs updating is performing calculations. So, jModelTest and bModelTest can, but do not necessarily agree on the most appropriate model to use.

## Conclusions

bModelTest is a BEAST 2 package which can be used in any analysis where trees are estimated based on nucleotide sequences, such as multi-species coalescent analysis [28, 29], various forms of phylogeographical analyses, sampled ancestor analysis [30], demographic reconstruction using coalescent [31], birth death skyline analysis [32], *et cetera*. The GUI support provided through BEAUti makes it easy to set up an analysis with the bModelTest site model: just select bModelTest instead of the default gamma site model from the combo box in the site model panel.

bModelTest allows estimation of the site model using a full Bayesian approach, without the need to rely on non-Bayesian tools for selecting the site model.

## List of abbreviations

BEAST: Bayesian evolutionary analysis by sampling trees
GTR: general time reversible
GUI: graphical user interface
HCV: hepatitis C virus
HPD: highest probability density
LGPL: Lesser General Public License
MCMC: Markov chain Monte Carlo

## Ethics approval and consent to participate

Not applicable.

## Consent for publication

The authors declare that they consent to pubication.

## Availability of data and material

See supplementary information for material for the simulation study as well as the primates and HCV analyses.

## Competing interests

The authors declare that they have no competing interests.

## Author’s contributions

RRB designed and developed software. AJD and RRB designed experiments. RRB executed experiments. AJD and RRB analyzed the data. AJD and RRB prepared the figures. AJD and RRB wrote the manuscript.

## Acknowledgements

Not applicable.

## Funding

This work was funded by a Rutherford fellowship (**http://www.royalsociety.org.nz/programmes/funds/rutherford-discovery/**) from the Royal Society of New Zealand awarded to Prof. Alexei Drummond.

## Additional Files

Appendix.pdf

List of all models and their dependencies

List of parameters reported by the model

95% HPD of models for Primates data

Table S1-S7 tables with simulation study results

~~~
simstudy.tgz
Archive containing XML input files for simulation study, and summaries of log files. Use tar fxz simstudy.tgz to
uncompress this archive. Once uncompressed, the following files become available:
simstudy/equalFreqGammInv/analysis-out0.xml ... analysis-out99.xml XML input files for data generated with equal
frequencies, gamma rate heterogeneity and a proportion invariable sites
simstudy/equalFreqGamm/analysis-out0.xml ... analysis-out99.xml XML input files for data generated with equal
frequencies, gamma rate heterogeneity and no proportion invariable sites
simstudy/equalFreqInv/analysis-out0.xml ... analysis-out99.xml XML input files for data generated with equal
frequencies, no gamma rate heterogeneity and a proportion invariable sites
simstudy/equalFreq/analysis-out0.xml ... analysis-out99.xml XML input files for data generated with equal
frequencies, no gamma rate heterogeneity and no proportion invariable sites
simstudy/unequalFreqGammInv/analysis-out0.xml ... analysis-out99.xml XML input files for data generated with
unequal frequencies, gamma rate heterogeneity and a proportion invariable sites
simstudy/unequalFreqGamm/analysis-out0.xml ... analysis-out99.xml XML input files for data generated with
unequal frequencies, gamma rate heterogeneity and no proportion invariable sites
simstudy/unequalFreqInv/analysis-out0.xml ... analysis-out99.xml XML input files for data generated with unequal
frequencies, no gamma rate heterogeneity and a proportion invariable sites
simstudy/unequalFreq/analysis-out0.xml ... analysis-out99.xml XML input files for data generated with unequal
frequencies, no gamma rate heterogeneity and no proportion invariable sites
simstudy/*/out/summary contains log-analyser summary of the output for a run of the XML files in its parent
directory.
simstudy/*/out/coverage contains coverage summary of the output for a run of the XML files in its parent directory.
simstudy/*/out/run.sh shell script to run XML files in its parent directory. Assumes that `runbeast.sh' is a script in
the path that starts BEAST 2 and that the bModelTest package installed.
simstudy/truth.dat table with parameters of site models used to generate sequence data. Note only those parameters relevant for a particular site model are used, and the others are ignored if they are not relevant. For example, for cases where equal frequencies are used to simulate the data, the frequency entries in truth.dat are ignored.
~~~

The log files are quite large (about 4 Gigabyte) and since the summary files are as informative they are not included in the available data. However, they can be reconstructed using the run.sh scripts.

In addition, the following files are available from https://github.com/BEAST2-Dev/bModelTest/releases/tag/simstudy

simstudy5taxa.tgz

As simstudy.tgz but with 5 taxa, relaxed clock and sampled trees.

simstudy5taxaHKY.tgz

As simstudy5taxa.tgz but with substitution model fixed to HKY.

simstudy5taxaGTRtgz

As simstudy5taxa.tgz but with substitution model fixed to GTR.

simstudy5taxaAll.tgz

As simstudy5taxa.tgz but with substitution models sampled from all reversible models. Contains fasta alignments for jModelTest and jModelTest output.

simstudy16taxa1k.tgz

As simstudy16taxa.tgz but with 16 taxa and sequences of 1K length.

simstudy16taxa0.5k.tgz

As simstudy16taxa1k.tgz but with sequences of 0.5K length.

hcv1.xml

XML input file for the hepatitis analysis. Requires BEAST 2 and the bModelTest package to run.

hcv1.log

Can be inspected using Tracer (available from http://tree.bio.ed.ac.uk/software/tracer/);

It can be analysed using the BModelAnalyser utility that package, by starting BEAUti (the GUI that comes with BEAST), select File/Launch Apps, then select BModelAnalyser from the list of available utilities. Select this log file in the dialog that shows up and a report with coverage statistics and a graph of the models is shown.

primates.xml

XML input file for the primate analysis using empirical frequencies. Requires BEAST 2 and the bModelTest package to run.

Primates.log

Log files of a run of primates.xml. Can be analysed like hcv1.log.

primatesE.xml

XML input file for the primate analysis using estimated frequencies. Requires BEAST 2 and the bModelTest

package to run.

PrimatesE.log

Log files of a run of primatesE.xml. Can be analysed like hcv1.log.

## Footnotes

[1] Estimated shape parameters only take values of the shape parameter in account in the portion of the posterior sample where gamma rate heterogeneity indicator is 1.

[2] The estimated proportion of invariable sites only take values of the parameter in account in the posterior sample where the invariant category was present.

## References

1. Posada, D., Crandall, K.A.: Modeltest: testing the model of dna substitution. Bioinformatics 14(9), 817–818 (1998)

2. Posada, D.: jModelTest: phylogenetic model averaging. Molecular biology and evolution 25(7), 1253–1256 (2008)

3. Darriba, D., Taboada, G.L., Doallo, R., Posada, D.: jModelTest 2: more models, new heuristics and parallel computing. Nature methods 9(8), 772–772 (2012)

4. Yang, Z.: Maximum likelihood phylogenetic estimation from DNA sequences with variable rates over sites: Approximate methods. Journal of Molecular Evolution 39(3), 306–314 (1994). doi:10.1007/BF00160154

5. Gu, X., Fu, Y.X., Li, W.H.: Maximum likelihood estimation of the heterogeneity of substitution rate among nucleotide sites. Mol Biol Evol 12(4), 546–57 (1995)

6. Waddell, P., Penny, D.: Evolutionary trees of apes and humans from DNA sequences. In: Lock, A.J., Peters, C.R. (eds.) Handbook of Symbolic Evolution, pp. 53–73. Clarendon Press, Oxford., ??? (1996)

7. Huelsenbeck, J.P., Larget, B., Alfaro, M.E.: Bayesian phylogenetic model selection using reversible jump markov chain monte carlo. Molecular Biology and Evolution 21(6), 1123–1133 (2004). doi:10.1093/molbev/msh123

8. Bouckaert, R.R., Alvarado-Mora, M., Rebello Pinho, J.a.: Evolutionary rates and hbv: issues of rate estimation with bayesian molecular methods. Antiviral therapy (2013)

9. Drummond, A.J., Bouckaert, R.R.: Bayesian Evolutionary Analysis with BEAST. Cambridge University Press, Cambridge (2015)

10. Wu, C.-H., Suchard, M.A., Drummond, A.J.: Bayesian selection of nucleotide substitution models and their site assignments. Mol Biol Evol 30(3), 669–88 (2013). doi:10.1093/molbev/mss258

11. Bouckaert, R.R., Heled, J., Kuhnert, D., Vaughan, T., Wu, C.-H., Xie, D., Suchard, M.A., Rambaut, A., Drummond, A.J.: BEAST 2: a software platform for bayesian evolutionary analysis. PLoS Comput Biol 10(4), 1003537 (2014). doi:10.1371/journal.pcbi.1003537

12. Tavare, S.: Some probabilistic and statistical problems in the analysis of dna sequences. Lectures on mathematics in the life sciences 17, 57–86 (1986)

13. Hasegawa, M., Kishino, H., Yano, T.: Dating the human-ape splitting by a molecular clock of mitochondrial dna. Journal of Molecular Evolution 22, 160–174 (1985)

14. Pereira, L., Freitas, F., Fernandes, V., Pereira, J.B., Costa, M.D., Costa, S., Maximo, V., Macaulay, V., Rocha, R., Samuels, D.C.: The diversity present in 5140 human mitochondrial genomes. Am J Hum Genet 84(5), 628–40 (2009). doi:10.1016/j.ajhg.2009.04.013

15. Rosenberg, N.A.: The shapes of neutral gene genealogies in two species: probabilities of monophyly, paraphyly and polyphyly in a coalescent model. Evolution (2003)

16. Jukes, T., Cantor, C.: Evolution of protein molecules. In: Munro, H.N. (ed.) Mammaliam Protein Metabolism, pp. 21–132. Academic Press, New York (1969)

17. Felsenstein, J.: Evolutionary trees from DNA sequences: a maximum likelihood approach. Journal of Molecular Evolution 17, 368–376 (1981)

18. Tamura, K., Nei, M.: Estimation of the number of nucleotide substitutions in the control region of mitochondrial DNA in humans and chimpanzees. Molecular Biology and Evolution 10, 512–526 (1993)

19. Kimura, M.: Estimation of evolutionary distances between homologous nucleotide sequences. Proceedings of the National Academy of Sciences 78(1), 454–458 (1981)

20. Posada, D.: Using MODELTEST and PAUP* to select a model of nucleotide substitution. Current protocols in bioinformatics, 6–5 (2003)

21. Green, P.J.: Reversible jump markov chain monte carlo computation and bayesian model determination. Biometrika 82, 711–732 (1995)

22. Pagel, M., Meade, A.: Bayesian analysis of correlated evolution of discrete characters by reversible-jump markov chain monte carlo. The American Naturalist 167(6), 808–825 (2006)

23. Bouckaert, R.R.: BEASTShell - scripting for bayesian hierarchical clustering. Submitted (2015)

24. Dawid, A.P.: The well-calibrated bayesian. Journal of the American Statistical Association 77(379), 605–610 (1982)

25. Drummond, A.J., Ho, S.Y.W., Phillips, M.J., Rambaut, A.: Relaxed phylogenetics and dating with confidence. PLoS Biol 4(5), 88 (2006). doi:10.1371/journal.pbio.0040088

26. Hayasaka, K., Gojobori, T., Horai, S.: Molecular phylogeny and evolution of primate mitochondrial dna. Molecular Biology and Evolution 5(6), 626–644 (1988)

27. Gray, R.R., Parker, J., Lemey, P., Salemi, M., Katzourakis, A., Pybus, O.G.: The mode and tempo of hepatitis c virus evolution within and among hosts. BMC evolutionary biology 11(1), 131 (2011)

28. Heled, J., Drummond, A.J.: Bayesian inference of species trees from multilocus data. Mol Biol Evol 27(3), 570–80 (2010). doi:10.1093/molbev/msp274

29. Ogilvie, H.A., Heled, J., Xie, D., Drummond, A.J.: Computational performance and statistical accuracy of *beast and comparisons with other methods. Syst Biol 65(3), 381–96 (2016). doi:10.1093/sysbio/syv118

30. Gavryushkina, A., Welch, D., Stadler, T., Drummond, A.J.: Bayesian inference of sampled ancestor trees for epidemiology and fossil calibration. PLoS Comput Biol 10(12), 1003919 (2014). doi:10.1371/journal.pcbi.1003919

31. Heled, J., Drummond, A.J.: Bayesian inference of population size history from multiple loci. BMC Evol Biol 8, 289 (2008). doi:10.1186/1471-2148-8-289

32. Stadler, T., Kuhnert, D., Bonhoeffer, S., Drummond, A.J.: Birth-death skyline plot reveals temporal changes of epidemic spread in hiv and hepatitis c virus (hcv). Proc Natl Acad Sci USA 110(1), 228–33 (2013). doi:10.1073/pnas.1207965110

